# The anticancer chemotherapy drug 5-Fluorouracil has positive interaction with antibiotics and can select for antibiotic resistance in *Staphylococcus aureus*

**DOI:** 10.1101/2023.09.25.559397

**Authors:** Sara R. Henderson, Leila H. Aras, Benjamin A. Evans

**Affiliations:** Norwich Medical School, University of East Anglia, Norwich, UK; Centre for Skin Sciences, University of Bradford, Bradford, UK

## Abstract

Antibiotic resistance is a major global issue in healthcare and understanding the drivers of resistance is key in developing effective strategies to counter it. Many non-antibiotic drugs, such as cancer chemotherapy drugs, can have antimicrobial properties but their effects on bacteria in the context of infection and drug resistance have only recently begun to be explored. Here we investigate the antimicrobial properties of the cancer drug 5-fluorouracil (5-FU) on the common human commensal and pathogen *Staphylococcus aureus*. 5-FU can be metabolized by *S. aureus* and ultimately results in the inhibition of ThyA, involved in the folate synthesis and thymine synthesis pathways. Bacterial growth was inhibited by 5-FU, and the drug had additive or synergistic interactions with the antibiotics trimethoprim and sulfamethoxazole. The addition of thymidine overcame the inhibitory effects of 5-FU. Strains of *S. aureus* evolved in the presence of 5-FU developed mutations in the thymidine kinase gene *tdk*, likely inhibiting the thymine salvage pathway. In mixtures of clinical trimethoprim-resistant *S. aureus* strains and sensitive strains, the presence of 5-FU conferred a large fitness advantage to the resistant strains and selected for them over the sensitive strains. Together these data show that 5-FU has antimicrobial effects against *S. aureus* with these effects targeting the same pathway as existing antibiotics, and that the use of 5-FU in patients may be selecting for antibiotic-resistant bacteria.

## Introduction

The increase in antibiotic resistance is one of the 13 biggest threats to global health, with the potential to reverse advancements in modern medicine including global cancer survival rates (1, 2). Since the introduction of antibiotics in the 1940s there has been clinical development of resistance, limiting treatment options. The situation we currently find ourselves in is concerning as antibiotic resistance is developing and spreading at a rate faster than we can generate new or optimized treatment options (3). Understanding how antibiotic resistance evolves and is selected is crucial for developing strategies to counter this threat. Cancer patients in particular are at risk of bacterial infection due to cancer- or treatment-induced reduced immune function, so effective antibiotic therapy is essential for managing these patients. Increasingly, there is recognition of a link between bacteria and cancer. Recent work has shown that bacteria alter the efficacy of chemotherapy drugs (4); in some cases bacteria are required for chemotherapy drugs to work effectively (5), while in others intra-tumour bacteria are antagonistic to chemotherapy drug action (6). While there is an emerging field of work directed at understanding how bacteria affect cancer development and chemotherapy effectiveness, the study of how chemotherapy drugs affect bacteria themselves has been neglected.

5-FU like many other anticancer-chemotherapy treatments was developed in the 1950s with initial indication of antibiotic properties (7), but it was further developed for its anticancer properties (8, 9). Although the drug has significant toxicities it is still routinely used in treatment programs including for skin, breast and colorectal cancers. Its use is highly effective due to its multiple modes of action including incorporation into DNA/RNA, metabolism to FUTP/FdUTP preventing RNA/DNA synthesis, and finally and possibly most importantly metabolism to FdUMP inhibiting the action of thymidylate synthase as reviewed by (10). The metabolism of 5-FU to FdUMP can occur via 2 major intermediates in humans, FUR or FdURD, each utilising 2 enzymes. However, in the bacterium *Staphylococcus aureus* the uridine phosphorylase enzymes (Upase) do not have bacterial homologues, hence this pathway is not present in bacteria. Both Pyrimidine/Thymidine phosphorylase (DeoA) and thymidine kinase (TDK) are both found in the bacterial kingdom. One key point to note is that bacterial TDKs cluster into 2 evolutionary groups with most Gram positive bacteria having a highly conserved TDK enzyme in comparison to the eukaryotic enzyme, whereas Gram negative species tend to have a greater degree of divergence and loss of conserved residues (11).

In humans there is a key pathway by which uracil (and thereby implied) fluorouracil can be detoxified and excreted through catabolism started by dihydropyrimidine dehydrogenase (12). This pathway has also been described for some bacterial organisms such as *Escherichia coli* (13) and *Clostridium uracilicum* (14–18), but no homologues have thus far been reported in Gram positive organisms such as *S. aureus*. This means that bacteria such as *S. aureus* are more susceptible to the effects of 5-FU as it is unable to directly detoxify 5-FU, hence making the inhibition of thymidylate synthase more important. Thymidylate synthase is universally conserved across all organisms performing the essential process of converting dUMP to dTMP for formation of thymidylate, while utilizing Tetrahydrofolate reforming dihydrofolate. This process links the folate cycle to the creation of thymidine for DNA synthesis. This is relevant in respect of antimicrobial resistance as the bacterial folate cycle is inhibited by the clinically used antibiotics sulfamethoxazole and trimethoprim.

Sporadic studies over the last 50 years have demonstrated that some cancer chemotherapy drugs such as bleomycin, doxorubicin, and cisplatin are antibacterial (and indeed some were originally proposed as antibacterials (7)), induce the bacterial SOS response and can cause phenomena such as bacteriophage induction (8, 9, 19–21). It is therefore apparent that anti-cancer drugs have profound effects on bacteria. An area of particular concern is the effect that chemotherapy drugs have on bacterial susceptibility to antibiotics, which is supported by two major lines of evidence. Firstly, when exposed to antibacterial chemotherapy drugs, bacteria use intrinsic mechanisms (*e.g.* upregulation of efflux such as the AcrB pump) to survive exposure to the drugs (22, 23). These are the same mechanisms used to survive antibiotic exposure. Therefore, it is likely that exposure to chemotherapy drugs will result in bacteria becoming less susceptible to antibiotics. Secondly, some bacteria carry genes that confer resistance to chemotherapy drugs (*e.g.* bleomycin). These genes are often adjacent to antibiotic resistance genes on plasmids that can be transferred between bacteria (24). Therefore, it is likely that exposure to chemotherapy drugs will result in co-selection of antibiotic resistance genes and associated plasmids.

Here, we investigate the impacts of 5-FU on the common human-associated bacterium *S. aureus*, with a particular focus on the antimicrobial trimethoprim and sulfamethoxazole. We find that 5-FU is inhibitory to *S. aureus*, and positively interacts with antimicrobials. Further, 5-FU can select for antibiotic-resistant strains of *S. aureus*.

## Methods

### Bacterial Strains

*S. aureus* ATCC 25923, NCTC 6571 and F77 were grown in Mueller-Hinton (MH) broth or agar unless otherwise stated. *S. aureus* strains 16, 42 and 60 are clinical isolates collected between 2015-2016 from hospitals in Alexandria, Egypt (25). Isolates were confirmed as *S. aureus* by genome sequencing. Trimethoprim, sulfamethoxazole and 5-FU were all solubilised in DMSO with concentrations kept below 2%.

### Experimental Evolution

*S. aureus* was grown for an overnight culture at 37°C and 180 rpm. A 1/100 dilution of overnight culture was made in final volume 100 µl MH broth containing 0, 0.025 or 6 µg/mL 5-FU and 1% (v/v) DMSO. Sequential passages were performed by performing a 1/100 dilution of the previous culture to fresh broth containing the appropriate concentration of 5-FU and DMSO. The experiment was conducted for a total of 15 passages.

### Minimum Inhibition Concentration testing

Broth MIC values were conducted as per the EUCAST guidelines, using the broth micro-dilution method in Mueller-Hinton broth (26). Checkerboard assays were performed in the same way as broth MIC assays except that two drug dilution series were used per plate, running perpendicular to one another, with the inoculum generated by colony resuspension (27). Disk diffusion MICs were conducted as per the EUCAST guidelines using 20 mL Mueller-Hinton agar plates and recommended disk concentrations (26). For 5-FU disks, 30 µg of 5-FU was added to a blank disk. All measurements were read manually using a ruler.

### Growth curves

Individual colonies of strains were grown for 8 hours at 37 °C shaking. These starter cultures were then diluted to a starting OD_600_ of 0.02 with a minimum of 3 repeats per condition. Growth was monitored in an *Omega FluroStar* plate reader at OD_600_ every 2-5 minutes for a minimum of 16 hours. Statistical analyses were performed using repeated measures ANOVAs with post-hoc Bonferroni testing in SPSS.

### Competition assays

Overnight cultures of *S. aureus* were adjusted to OD_600_ of 0.1. Adjusted cultures were diluted 10-fold in MHB to give an inoculum of ∼10^6^ CFU/mL and then mixed in a 1:1 ratio in a new 5 mL tube. A 50 µL volume of the strain mixture was added to each of six tubes containing 5 mL MHB and vortexed. Immediately, 50 µL from each tube was removed and plated out on MHA and MHA + 100 µg/mL TMP at dilutions of 10^1^, 10^2^, and 10^3^ in duplicate. The plates were then incubated overnight at 37° C. The colony counts from these plates were used to calculate the initial frequencies of the strains in the mixture. As soon as the 50 µL was removed, 5-FU was added to three of the tubes to a final concentration of 30 µg/mL. All tubes were incubated overnight at 37° C with orbital shaking at 180 rpm. Following incubation, 50 µL from each of the six tubes was plated on MHA at dilutions between 10^3^ – 10^6^ in duplicate, and on MHA + 100 µg/mL TMP at dilutions between 10^4^ – 10^7^ in duplicate. Colony counts from these plates were used to calculate the number of sensitive and resistant strains following competition. Relative fitness of each strain in the selection experiments was calculated using the equation below (28):

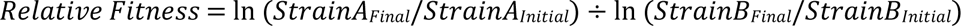

where strain A and strain B are TMP resistant and sensitive respectively, and ‘ln’ is the natural logarithm. Subscripts ‘initial’ and ‘final’ indicate the numbers of colonies before and after the selection experiment. Differences between treatments were determined using 2-tailed t-tests.

### DNA extraction and sequencing

For Illumina sequencing bacteria were lysed by bead beating using Beadbug 0.1 mm silica bead tubes (Sigma-Aldrich) and beaten using a Qiagen Tissue Lyser II at a frequency of 30 hertz for 3 minutes. Genomic DNA was then extracted and purified using the Wizard Genomic DNA kit (Promega, UK). Purified DNA was Illumina sequenced at the Quadram Institute, Norwich, UK. The ancestral strains were assembled and annotated to form reference genomes using a pipeline of Trimmomatic (29), SPAdes (30), and Prokka (31). Assemblies were checked for completeness and contamination using CheckM (32). Polymorphisms between the evolved and ancestral strains were detected using Breseq (33). Selected single nucleotide polymorphisms detected were then PCR amplified and sent for Sanger sequencing to confirm the presence of the SNP.

## Results

### Staphylococcus aureus has the enzymes to metabolise 5-FU

In humans there are several enzymatic pathways by which uracil is metabolised. The 5-fluoro analogue 5-FU can also be metabolised by many of these enzymes. The major pathways of interest are those that result in FdUMP which is a fluoro analogue of dUMP which in the fluorinated form inhibits the essential enzyme Thymidine synthase (TS/ThyA). The two major pathways in humans for this process are via TP and TK1 or through Upase1/2 and UCK1/2 (Figure 1). In *S. aureus* there are no homologues of Upase1/2 meaning that this pathway is not present. On the other hand, there are homologues of TP (DeoA) and of TK1 (Tdk). Therefore, we postulated that 5-FU was likely to exhibit antimicrobial effects on *S. aureus* through metabolism by DeoA and Tdk to FdUMP resulting in inhibition of the essential enzyme ThyA. In anticancer chemotherapy treatment 5-FU is co-administered with folinic acid to increase the activity of the folate cycle which is inhibited by the action of Trimethoprim and sulphamethoxazole, hence we also theorised that there might be interactions (antagonism) between the folate inhibiting antimicrobials and 5-FU.

**Figure 1:**
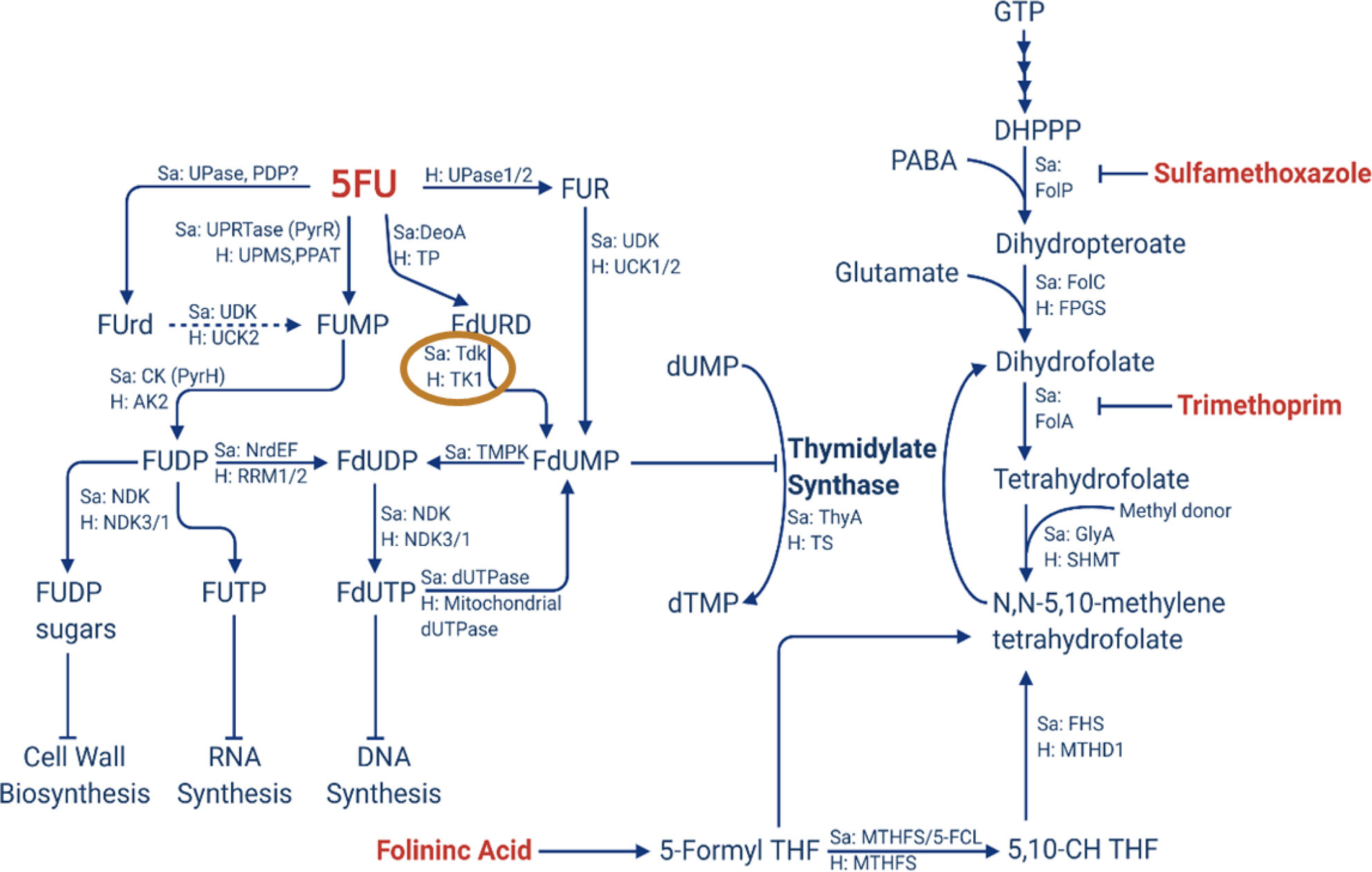
Metabolism and mechanism of action of 5-fluorouracil, and it’s relationship to the folate synthesis pathway. The inhibitory drugs 5FU, trimethoprim and sulfamethoxazole are shown in red. Alongside arrows, enzymes found in *S. aureus* are indicated by ‘Sa’, while those found in humans are indicated by ‘H’. The enzymatic step effected by mutation in the current study is circled in gold.

### 5-Fluorouracil is an antimicrobial to *S. aureus*

To confirm that 5-FU had antimicrobial effects on *S. aureus* we initially performed MIC tests on *S. aureus.* We also tested our strains for susceptibility to trimethoprim, sulphamethoxazole and co-trimoxazole. The strain F77 was found to be clinically resistant to trimethoprim, sulphamethoxazole, cotrimoxazole (potentially due to the carriage of *dfrC* (34)), and also had decreased susceptibility to 5-FU (Table 1). In contrast, strain NCTC 6571 was susceptible to all of these antibiotics with increased sensitivity to 5-FU (Table 1). This suggested that there may be some overlap of the resistance mechanisms for these compounds.

**Table 1:**
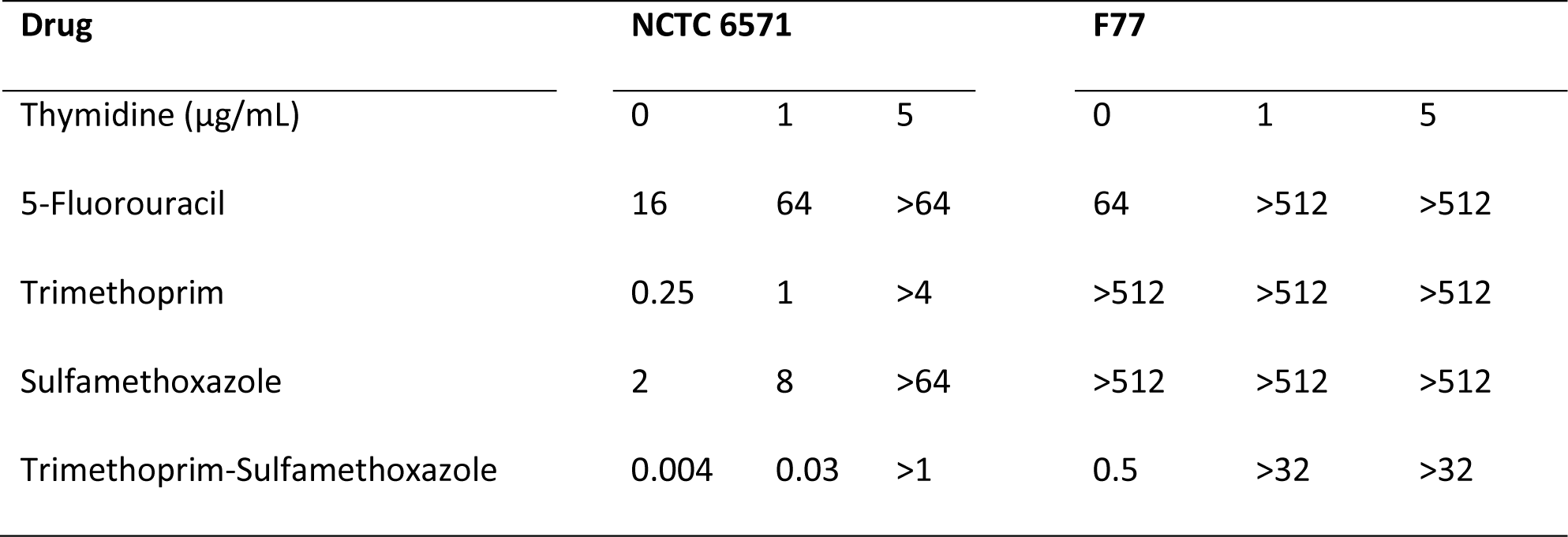
MIC values (µg/mL) for the ancestral *S. aureus* NCTC 6571 and F77 strains in the presence and absence of thymidine.

### 5-FU has some interaction with folate inhibiting antibiotics

We were interested in how 5-FU interacted with trimethoprim and sulphathiazole. Initially we ran checkerboard assays to obtain the fractional inhibition concentration (FIC) indices. These showed the FIC indices to be between 0.5 – 1 indicating the interactions between the drugs are mostly synergistic or additive (Table 2) (35). To compliment these results, we ran growth curves in sub-MIC (0.5 x MIC) combinations of the same drugs (Figure 2). Our results showed that for both strains tested (NCTC 6571 and F77) there was no difference in growth compared to the control in the presence of TMP alone, and a small but significant difference in the presence of SFX alone for F77 (*p* = 8.6190 x 10^-5^). In the presence of sub-MIC concentrations of 5-FU the growth was significantly reduced compared to the control (NCTC 6571: *p* = 3.3166 x 10^-6^; F77: *p* = 6.6163 x 10^-19^). With the TMP/SFX combination, as expected, there was also a significant reduction in growth relative to the control (NCTC 6571: *p* = 0.001; F77: *p* = 5.6847 x 10^-20^) demonstrating the synergism between these two drugs. When TMP was added to 5-FU the growth profile was found to be significantly impaired and similar to that with just 5-FU alone. Of note was that when SFX was added to 5-FU, the growth profile was intermediate between that for 5-FU or SFX alone, with the strain growing significantly better than when 5-FU alone was used (NCTC 6571: *p* = 0.003; F77: *p* = 2.2368 x 10^-11^), and in the case of NCTC 6571 not differing from the positive control (NCTC 6571: *p* = 0.13). This suggest that this interaction between SFX and 5-FU may therefore be antagonistic at sub-MIC values (Figure 2).

**Figure 2:**
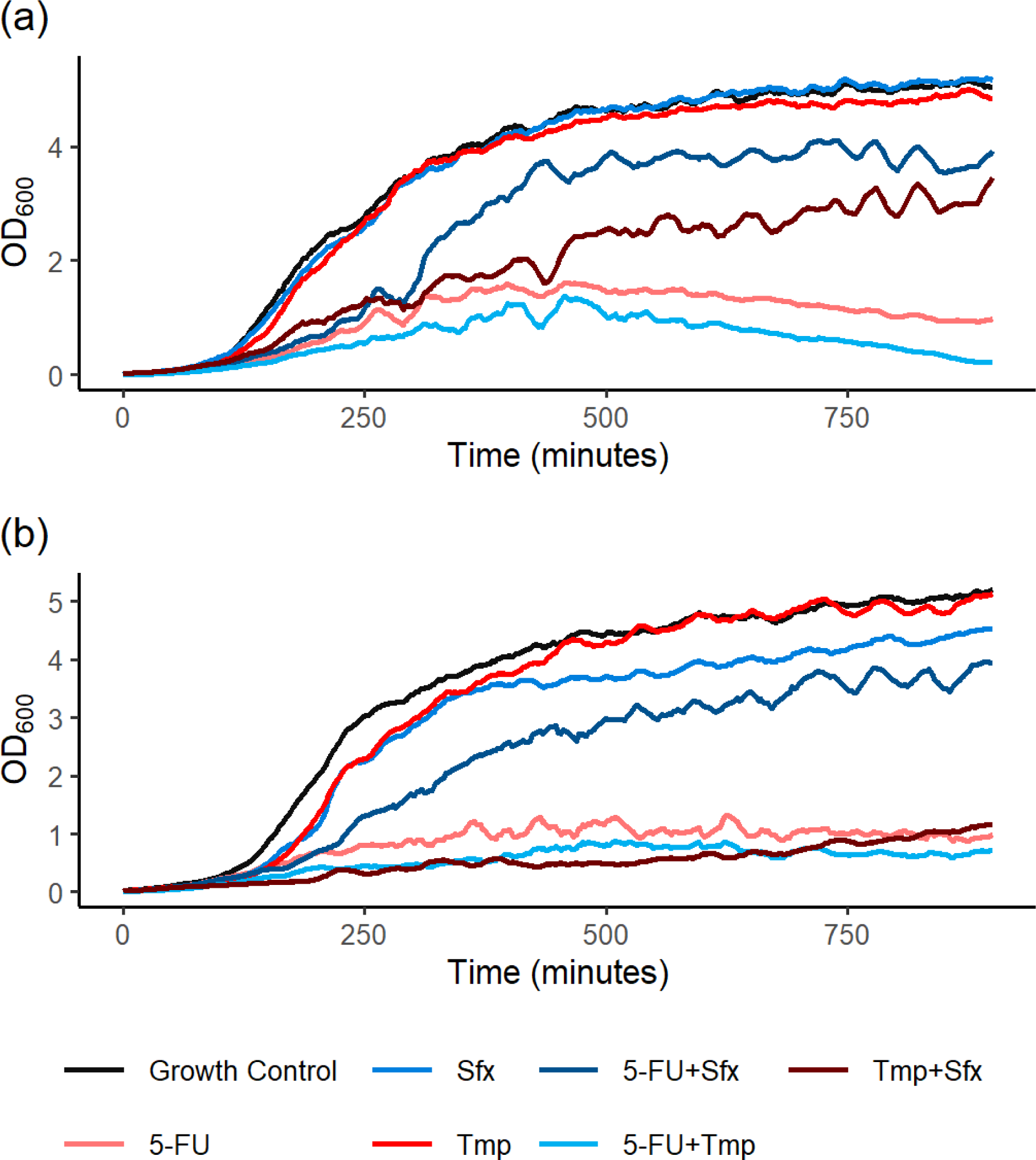
Growth of *S. aureus* in Mueller-Hinton broth a) in the presence of NCTC6571 and b) F77 in the presence of 5-FU a) 32 mg/L, b) 8 mg/L); Sulfamethoxazole a) 32 mg/L, b) 4 mg/L; Trimethoprim a) 32 mg/L and b) 0.25 mg/L. Growth curves are averages of minimum 4 technical replicates.

**Table 2:**
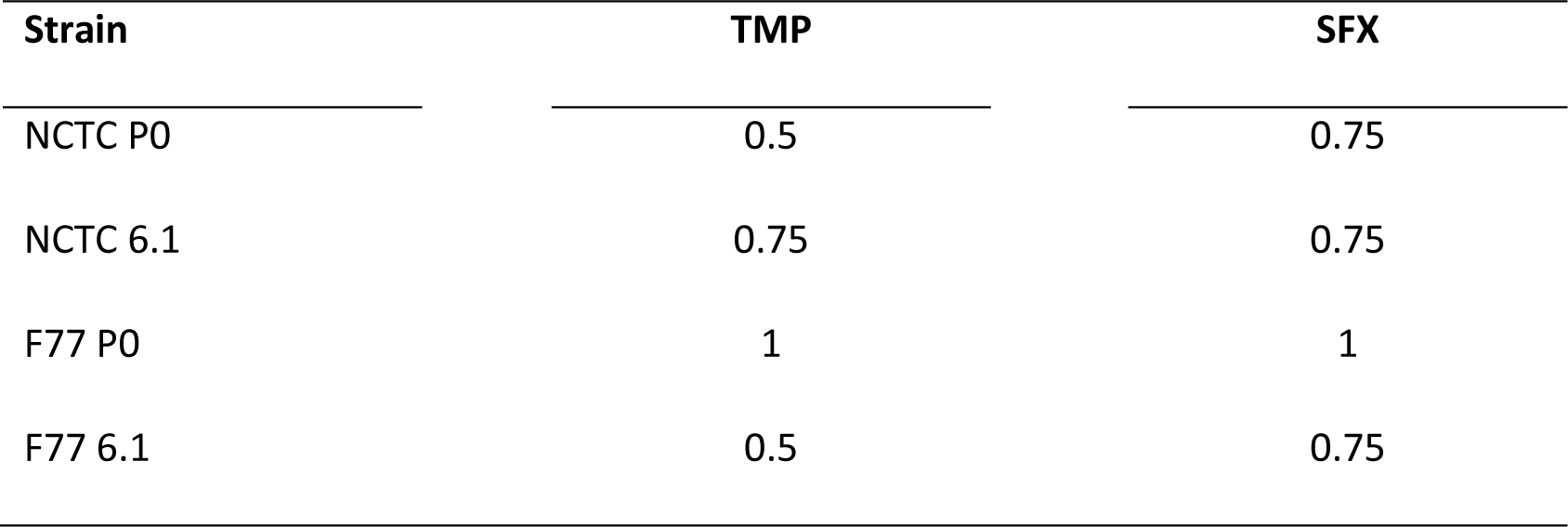
Fractional Inhibitory Concentration (FIC) indices of trimethoprim or sulfamethoxazole with 5-FU. Values ≤0.5 indicate synergism, >0.5 to 1 suggest an additive effect, >1 to <2 indicates indifference, and ≥2 indicates antagonism(35).

| Strain | TMP | SFX |
| --- | --- | --- |
| NCTC P0 | 0.5 | 0.75 |
| NCTC 6.1 | 0.75 | 0.75 |
| F77 P0 | 1 | 1 |
| F77 6.1 | 0.5 | 0.75 |

As folinic acid is co-prescribed with 5-FU in anticancer chemotherapy we were interested to understand how it alters the growth profile of *S. aureus* in the presence and absence of 5-FU and TMP. Our results showed that at physiologically relevant concentrations folinic acid does not influence growth in the presence or absence of 5-FU or TMP at sub-MIC values (Figure S1).

### Thymidine increases tolerance to 5-FU

It has been reported that in the presence of thymidine there is increased tolerance to trimethoprim, sulfamethoxazole and cotrimoxazole because it bypasses the thymine synthesis pathway and therefore the inhibitory effects of the drugs (36). Therefore, we were interested to know if increased tolerance to 5-FU was achieved in the presence of excess thymidine. We initially assessed MICs which showed a 4-fold increase in MIC value in the presence of just 1 mg/L thymidine (Table 1). We investigated this further through growth curves in the presence of thymidine. These showed that at concentrations of trimethoprim and 5-FU where growth was normally prevented, growth was supported in the presence of 5 mg/L thymidine (Figure 3). Together, these data show that, as with cotrimoxazole, the addition of thymidine overcomes the inhibitory effects of 5-FU, providing further evidence that cotrimoxazole and 5-FU inhibit growth through a related mechanism.

**Figure 3:**
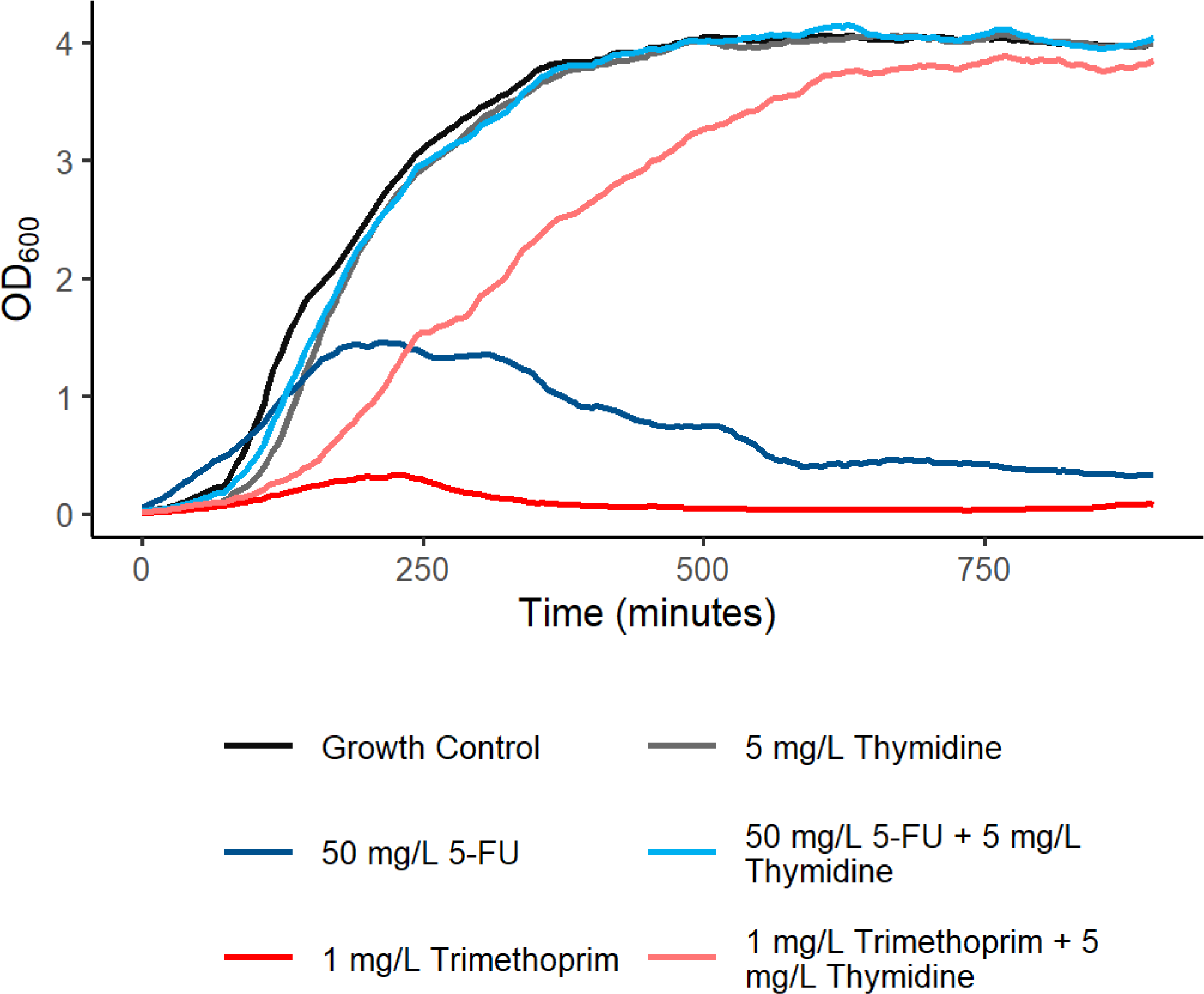
Effect of thymidine (5 mg/L) on the growth of *S. aureus* NCTC6571 in Mueller-Hinton broth in the presence or absence of 50 mg/L 5-FU and 1 mg/L trimethoprim. Growth curves are averages of minimum 4 technical replicates.

### Evolution of S. aureus in the presence of 5-FU results in resistance to 5-FU but not clinical antibiotics

Our data so far has shown the effects of acute exposure of strains to 5-FU. To compliment this we investigated how longer term exposure to the drug effects the bacteria. We passaged *S. aureus* ATCC 29213, NCTC 6571 and F77 strains 15 times, representing ∼200 generations, in the presence of 6 mg/L or 0.025 mg/L 5-FU or a solvent control (1% DMSO) in 4 technical replicates. Concentrations of 5-FU were chosen to be the maximal concentration that did not impede growth in the presence of 5-FU and the approximate concentration that might be in the body 4 hours post-treatment based on the biological half-life and C_max_ concentrations (37, 38). After 15 passages we performed disk diffusion assays on a panel of antimicrobial agents and 5-FU to investigate differences between the ancestral strains and the evolved strains. For strain NCTC 6571 we saw complete resistance to 5-FU with no region of inhibition being observed on application of 5-FU disks containing 30 mg/L (Table 3). The F77 strain that started with greater tolerance to 5-FU was unable to further adapt to 5-FU and no change in susceptibility was observed. The ATCC 25923 strain generated increased tolerance to 5-FU however it still had some susceptibility at high concentrations as seen with the zone diameter decreasing from 25 mm in the controls compared to 12 mm in the 5-FU evolved strains. Our results also showed that there was no observable adaptation to the panel of clinical and non-clinical antimicrobial agents when using this method (Table 3).

**Table 3:** Disk diffusion MIC values (diameter in mm) for strains evolved against either 5-FU or a solvent control vs the ancestral strains. Data are an average of 4 duplicate daughter strains with a minimum of 2 duplicate readings. AMP – Ampicillin, CHL – Chloramphenicol, CIP – Ciprofloxacin, FU – 5-Fluorouracil, GEN – Gentamycin, TET – Tetracycline, TMP – Trimethoprim, STX – Trimethoprim-Sulfamethoxazole.

| Strain | Timepoint and treatment | AMP | CHL | CIP | FU | GEN | TET | TMP | SXT |
| --- | --- | --- | --- | --- | --- | --- | --- | --- | --- |
| ATCC 25923 | Ancestor | 39 | 33 | 30 | 26 | 33 | 34 | 32 | 42 |
|  | Endpoint control | 41 | 33 | 32 | 25 | 31 | 35 | 31 | 40 |
|  | Endpoint 5FU 0.25 ug/ml | 41 | 34 | 32 | 24 | 34 | 35 | 32 | 41 |
|  | Endpoint 5FU 6 ug/ml | 41 | 33 | 32 | 12 | 33 | 35 | 32 | 40 |
| NCTC 6571 | Ancestor | 41 | 34 | 35 | 29 | 31 | 35 | 38 | 40 |
|  | Endpoint control | 43 | 33 | 36 | 32 | 31 | 36 | 37 | 40 |
|  | Endpoint 5FU 0.25 ug/ml | 43 | 35 | 39 | 25 | 33 | 38 | 38 | 41 |
|  | Endpoint 5FU 6 ug/ml | 42 | 34 | 37 | 0 | 30 | 37 | 40 | 40 |
| F77 | Ancestor | 15 | 33 | 31 | 19 | 32 | 34 | 0 | 29 |
|  | Endpoint control | 14 | 30 | 32 | 24 | 30 | 29 | 0 | 29 |
|  | Endpoint 5FU 0.25 ug/ml | 15 | 31 | 33 | 19 | 32 | 34 | 0 | 31 |
|  | Endpoint 5FU 6 ug/ml | 15 | 31 | 31 | 22 | 30 | 32 | 0 | 30 |

We performed growth curves to determine if the resistance evolved to 5-FU affected the growth kinetics of the bacteria. The evolved strains grew with equivalent efficiency to the parent strain when growth is performed in the absence of 5-FU. In fact, in the absence of 5-FU the evolved control and 5-FU exposed strains both appear to have a small degree of adaptation to the growth conditions demonstrated by slightly improved growth kinetics compared to the ancestor (control: *p* = 0.008; 5-FU exposed, *p* = 0.005) but did not differ from each other (*p* = 1) (Figure S2). Similarly, growth of the ancestor and the evolved control strains was inhibited to the same degree by 5-FU (*p* = 1), whereas the strains that evolved with 6 mg/L 5-FU grew equally as well when exposed to 5-FU as the ancestor did in the absence of drug (*p* = 1).

As efflux can be involved in drug resistance including to trimethoprim (39) we performed efflux assays with both trimethoprim and 5-FU. Our results demonstrated that there was no observable difference between the efflux rate when either no treatment, 0.25 x MIC 5-FU or 0.25 x MIC trimethoprim was added to the NCTC 6571 or F77 strains (Figure 4A, B). Likewise, the strains evolved in the presence of 5-FU did not have alter efflux compared to those evolved in the presence of a solvent control (Figure 4C, D). Therefore, efflux did not appear to contribute to the altered phenotype.

**Figure 4:**
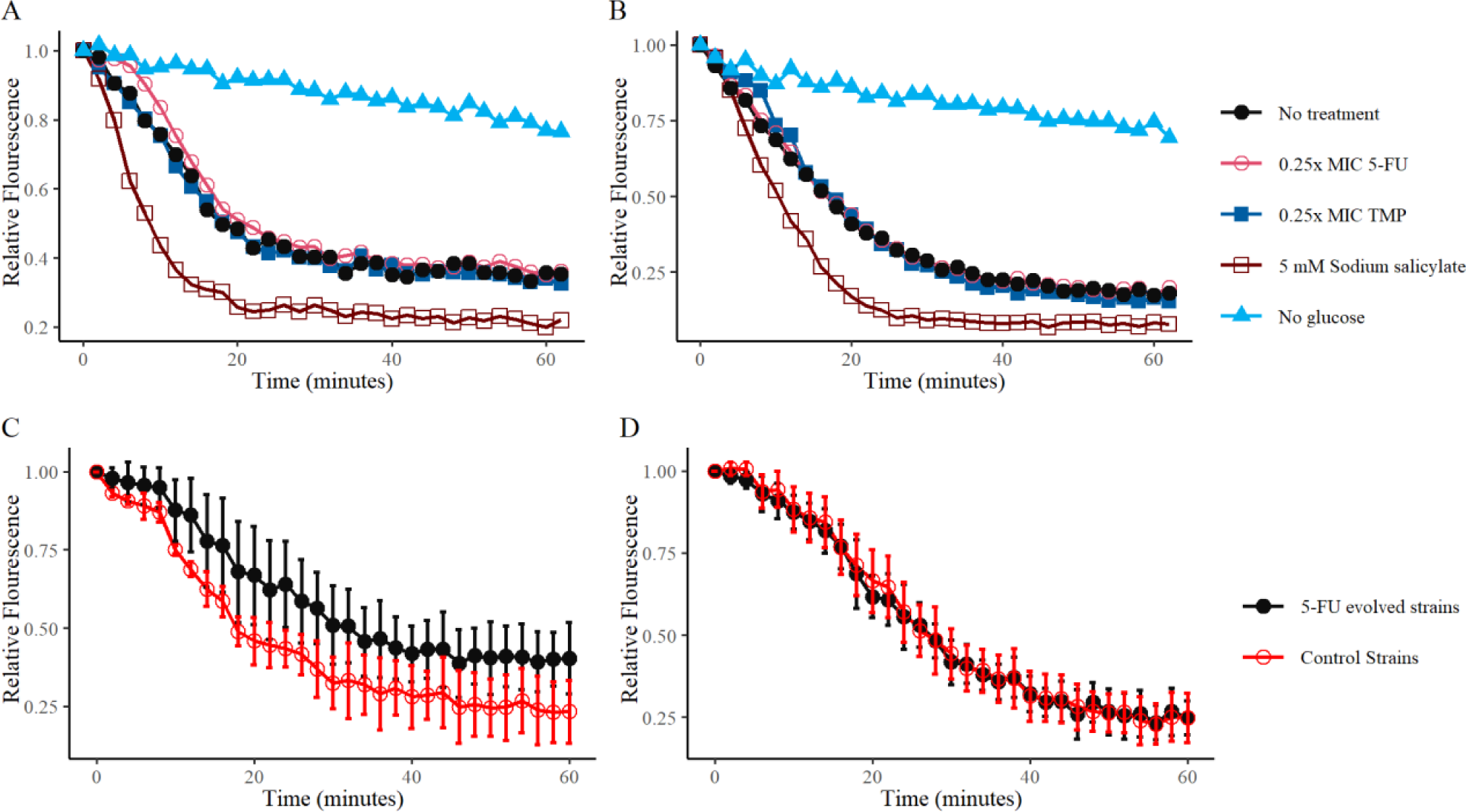
Efflux is not a resistance mechanism for *S. aureus* against 5-FU. Efflux of A) NCTC6571 or B) F77 in response to 0.25x MIC 5-FU or trimethoprim. C) Efflux of NCTC6571 or D) F77 strains evolved in the presence of 6 mg/L 5-FU or a solvent control. Error bars where present show the standard deviation of minimum 2 biological replicates or multiple technical replicates.

To understand the mechanism by which the resistance to 5-FU was obtained we performed Illumina whole genome sequencing on the ancestral and endpoint NCTC 6571 and F77 strains. We did not find any mutations that were consistent across the F77 strains that would suggest parallel evolution (Table 4). In contrast, in the NCTC 6571 daughter strains evolved with 5-FU, we observed mutations in the *tdk* gene, coding for thymidine kinase, in three out of the four daughter strains and no mutations in this region in the control strains. These mutations include two missense mutations and a premature stop codon resulting in non-functional *tdk* genes (Table 5). In the fourth daughter strain we observed a SNP 12 base pairs upstream of the *tdk* gene which is hypothesised to be within the Shine Dalgarno Region for this gene, potentially altering transcription of *tdk* (Table 5). All mutations were confirmed by sanger sequencing. While the precise impacts of these different mutations is unknown, the independent parallel evolution of different mutations in the *tdk* gene is strong evidence for selection at this locus.

**Table 4:**
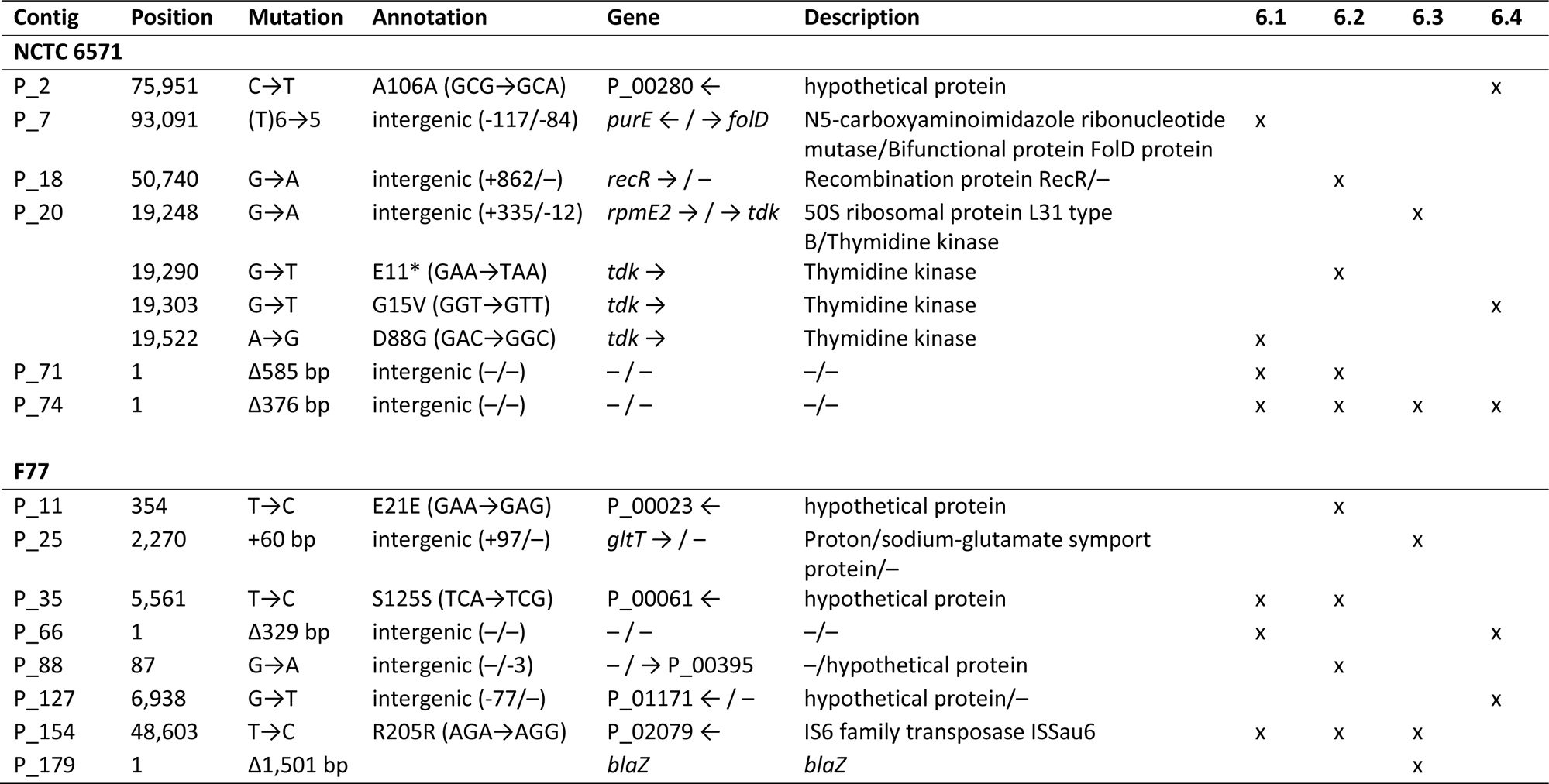
mutations unique to the strains evolved in 6 mg/L 5-FU compared with the control and mapped to the ancestral strain. The four strains evolved in the presence of 5-Fu are labelled 6.1, 6.2, 6.3 and 6.4.

### 5-FU can select for trimethoprim-resistant clinical strains

Due to the interactions between 5-FU and antibiotics that inhibit folic acid synthesis, we investigated whether 5-FU could select for antibiotic-resistant clinical strains. Three different clinical strains resistant to trimethoprim were competed against the antibiotic-sensitive type strain NCTC 6571. We found that all three clinical strains were slightly less fit than the sensitive strain when grown in the absence of any drug (Figure 5). However, when exposed to 5-FU all three clinical strain substantially outcompeted the sensitive strain, indicating that in a mixed population, 5-FU will select for pre-existing antibiotic-resistant strains.

**Figure 5:**
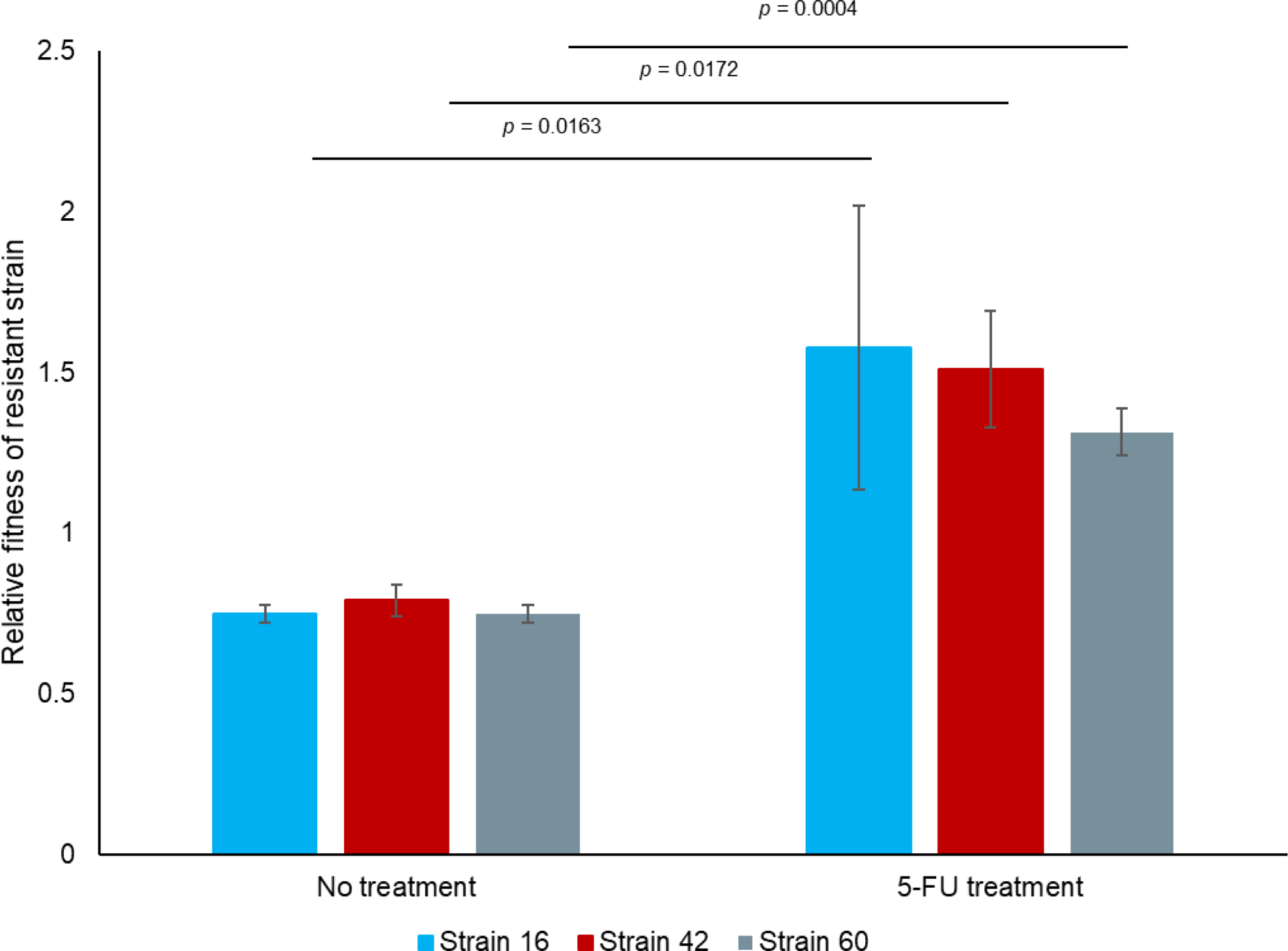
Relative fitness of 3 trimethoprim-resistant clinical isolates of *S. aureus* compared to the sensitive strain NCTC 6571 in the presence of no selection or in the presence of 30 µg/mL 5-FU. Data is from a minimum of 2 biological replicates.

## Discussion

Much progress has been made in recent years to better understand antimicrobial resistance evolution and spread. However, the vast majority of this work has focussed on drugs specifically prescribed for treating bacterial infections, such as antibiotics. Many other chemicals and drugs that bacteria may encounter in the healthcare setting are known to be antimicrobial, such as disinfectants or cancer chemotherapy drugs (20), yet little is known about how these other drugs might be affecting bacteria and whether there is an association with AMR. Here, we show that the cancer chemotherapy drug 5-FU inhibits bacterial growth via the same metabolic pathway as the antimicrobials trimethoprim and sulfamethoxazole, and can select for resistant strains.

It has been known for decades that some cancer chemotherapy drugs have antibacterial properties, including 5-FU. Here we show that 5-FU can significantly impair growth of *S. aureus* at concentrations that are found in patients receiving 5-FU treatment (40–42). Importantly, growth was affected at 5-FU concentrations that were below the MIC for the drug. Similar findings were recently published for the cancer chemotherapy drug methotrexate, which also acts through inhibiting the folate synthesis pathway (43). It is now appreciated that the sub-MIC selective window is important in the evolution of drug resistance, with concentrations of drug far below the MIC capable of selecting for the evolution of resistance over time (44). We observed this phenomenon here with *S. aureus* with the evolution of much greater tolerance to 5-FU after exposure to sub-inhibitory concentrations for ∼200 generations. Similar studies where increased tolerance to a non-antibiotic drug has evolved during experimental evolution have found corresponding evolution of cross-resistance to clinically used antimicrobials, such as trimethoprim in the case of methotrexate (43) or quinolones, ciprofloxacin and tetracycline in the case of disinfectants (45). We did not detect evidence for this for 5-FU, however the method used – disc diffusion – is not particularly sensitive and further analyses using dose response curves are required to confirm these results. However, we did find evidence for interactions between 5-FU and the antimicrobials trimethoprim and sulfamethoxazole. Checkerboard assays indicated that both the antimicrobials had synergistic or additive effects with 5-FU, consistent with their modes of action in inhibiting different parts of the folate synthesis pathway. This raises the possibility that in patients, the effectiveness of trimethoprim/sulfamethoxazole may be enhanced if 5-FU is also present, while on the other hand selection for reduced susceptibility by any one for the drugs could reduce the effectiveness of the others.

To try to understand the differences in phenotypic susceptibility to 5-FU we investigated the genetic changes in the evolved strains. Common to all strains that evolved increased tolerance to 5-FU were mutations in the coding region, or promotor region, of the *tdk* gene that encodes for thymidine kinase, which is involved in thymine salvage pathway (46). Nucleotide synthesis is important for bacterial pathogens (47). Knocking out the pyrimidine synthesis repressor gene *pyrR* results in increased pyrimidine synthesis and increased gut colonisation (48). Alternatively, inactivation of the purine synthesis repressor gene *purR* results in increased purine sythesis, leading to hypervirulence and drug resistance (49, 50). The role of nucleotide salvage pathways in bacterial pathogens is far less studied, and so it is not clear what the implications of inhibiting the thymine salvage pathway, through inactivation of *tdk*, may have on the wider bacterial phenotype. Inhibiting thymine salvage may result in the bacteria compensating by increasing *de novo* thymine sythesis. This could explain why we observed no growth defects in the strains carrying mutations in *tdk*. It is possible that these strains could have altered virulence, as seen with increased pyrimidine production, but this has not been tested here. Mutations in *thyA*, which catalyses the reaction in the thymine synthesis pathway that metabolised 5-FU products (namely fluorodeoxyuridylate – FdUMP) inhibit, are associated with the formation of small colony variants (SCVs) in *S. aureus* (52). However, the growth medium used here, Mueller Hinton broth and agar, is not thought to support the growth of *thyA* mutants (53), which may be the reason why mutations in *thyA* were not detected in this work. This is consistent with our strains mutating *tdk* instead – if the growth medium used doesn’t support the growth of *thyA* mutants due it not containing exogenous thymine that can be salvaged, it would support growth of mutants in the salvage pathway that can still *de novo* synthesise thymine. Furthermore, the *tdk* mutants isolated did not demonstrate a typical SCV phenotype, such as reduced growth and increased antimicrobial tolerance (52). This suggests that the overall impact of *tdk* mutations is quite different to *thyA* mutations, and that exposure to 5-FU in these conditions is not selecting for the evolution of a more generally antibiotic-resistant phenotype such as seen in SCVs.

The experiments conducted here were all performed in laboratory growth media, and therefore lack many features that bacteria would experience during human colonisation or infection. Work investigating the properties of *S. aureus* required for survival in human blood found that the pyrimidine salvage pathway is inhibited by human blood, and to be able to grow well the *S. aureus* need increased levels of pyrimidines (51). Therefore, in patients being treated with 5-FU, mutations in the pyrimidine salvage pathway to protect against 5-FU exposure may not carry as large a fitness cost if the bacteria are not relying as much on the salvage pathway in the first place. Furthermore, when the *S. aureus* are exposed to trimethoprim, that inhibits folate synthesis, the pyrimidine salvage pathway was required for growth. This would suggest that 5-FU and folate synthesis-inhibiting antimicrobials could work synergistically by inhibiting both synthesis and salvage of thymine, and is consistent with the results we present here. Therefore, despite the *in vitro* nature of our work, we believe that our observations are relevant to the clinical setting.

In addition to the potential for non-antibiotic drugs to promote the *de novo* evolution of antibiotic resistance, they can also select for pre-existing drug-resistant strains. Using three different clinical strains of *S. aureus* resistant to trimethoprim, we found that in the presence of 5-FU the clinical strains had a significant fitness advantage over the susceptible strain. It should be noted that it is not known whether it is due to the pre-existing resistance to trimethoprim held by the clinical strains that is responsible for this increased fitness in the presence of 5-FU, or whether it is some other aspect of these strains (for example, they were also resistant to a number of other antimicrobials including aminoglycosides, quinolones and linezolid (25)). Nevertheless, the ability of 5-FU to select for existing MDR *S. aureus* strains at sub-MIC concentrations is concerning as it suggests that in clinical usage 5-FU could promote the survival of resistant strains over their sensitive counterparts. This phenomenon is not restricted to 5-FU. Recent work has shown a similar effect with methotrexate, which selected for antibiotic resistant *E. coli* (43). The possibility that many non-antibiotic drugs commonly used in healthcare may be selecting for antimicrobial resistant bacteria is an area that merits further study if we are to fully understand antimicrobial resistance evolution and epidemiology and effectively combat this major healthcare issue.

## Acknowledgements

This work was funded by a Wellcome Trust Seed Award (grant reference 213979/Z/18/Z) awarded to BAE, and internal funding from the University of East Anglia. We are grateful to Lina Maarouf for providing us with clinical strains of *S. aureus*.

---

**Figure S1:**
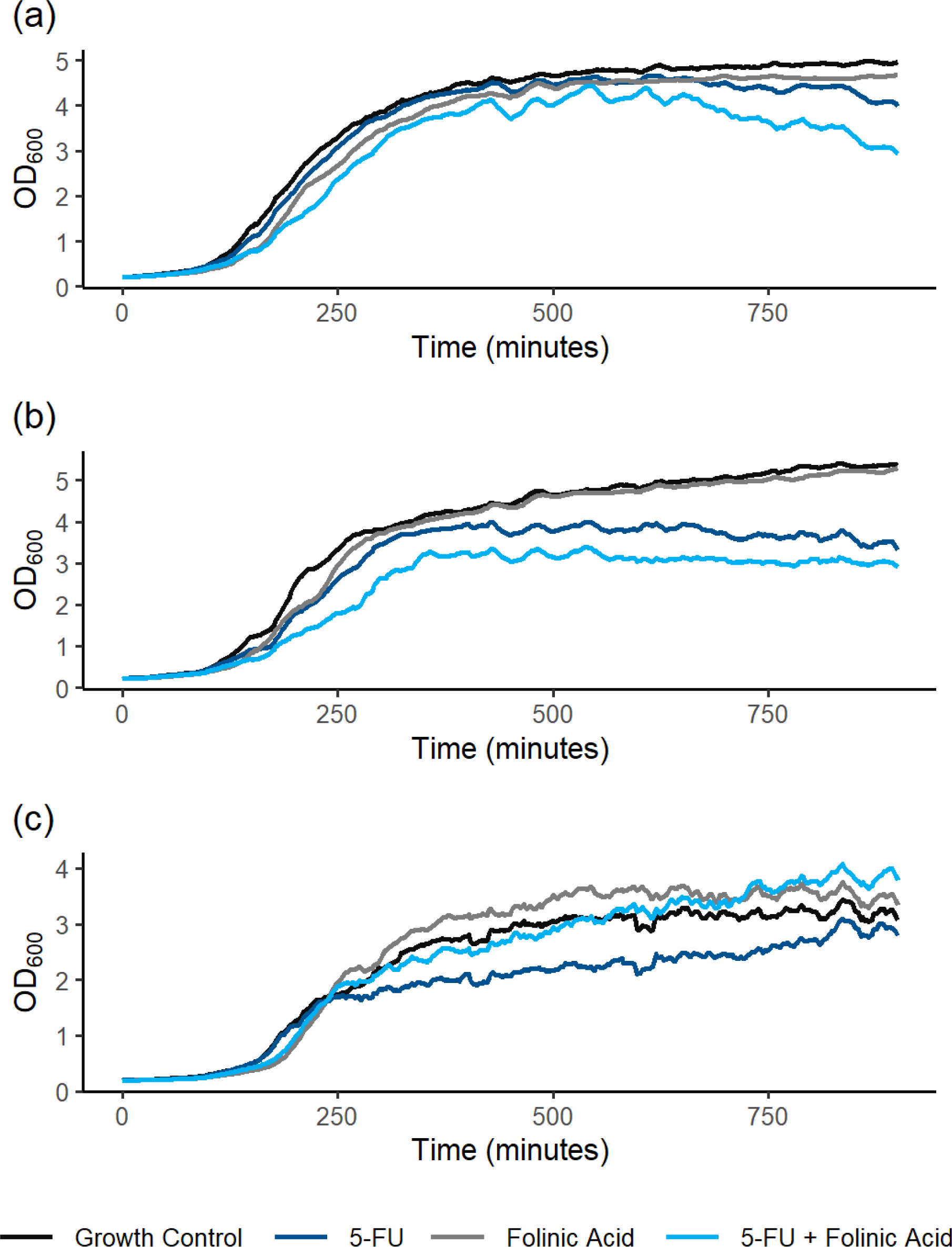
Folinic Acid does not impact on *S. aureus* growth. A) *S. aureus* NCTC6571, *S. aureus* ATCC25923 in the presence and absence of Folinic Acid (20 mg/L) and 5-FU F77 20 mg/L or NCTC6571/ATCC25923 5 mg/L.

**Figure S2:**
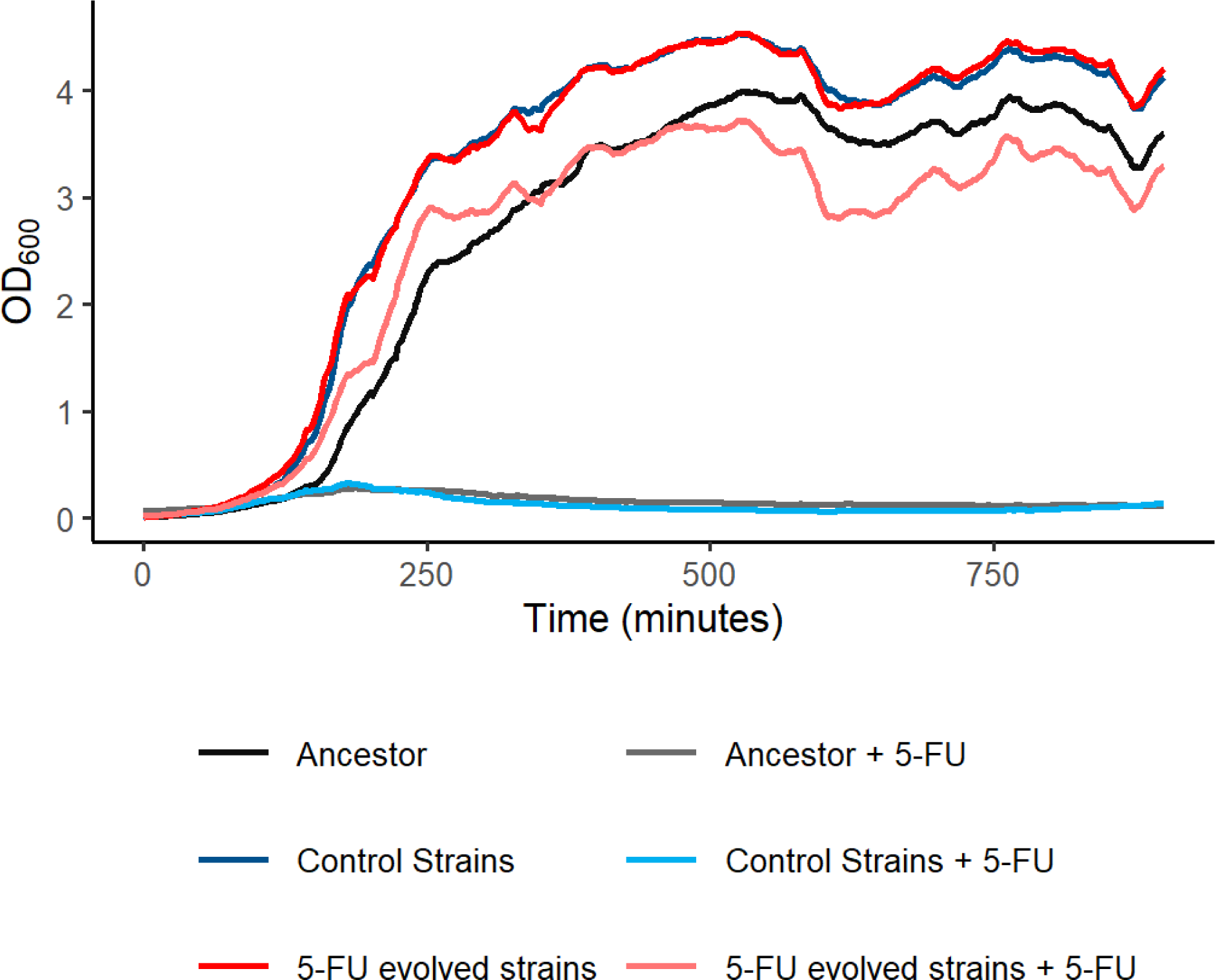
*S. aureus* NCTC6571 evolved in the presence of 6 mg/L 5-FU does not exhibit a fitness penalty but is able to grow in 16 mg/L 5-FU that previously it was unable to grow in. All growth curves were performed as the average of 4 biological replicate strains in Mueller-Hinton broth.

